# To share or not to share? iEEG evidence for state-dependent inequity encoding in the human OFC

**DOI:** 10.1101/2022.06.23.497432

**Authors:** Deborah Marciano, Brooke R. Staveland, Jack Lin, Ignacio Saez, Ming Hsu, Robert T. Knight

## Abstract

Social decision-making requires the integration of reward valuation and social cognition systems, both dependent on the orbitofrontal cortex (OFC). How these two OFC functions interact is largely unknown. We recorded intracranial activity from the OFC of ten patients making choices in the context of different types of inequity (disadvantageous vs. advantageous). We found that high-frequency activity (70-150 HZ) encoded the amount of self-reward, consistent with previous reports. We also observed novel evidence for encoding in human OFC of the social counterpart’s reward as well as the type of inequity being experienced. Additionally, we find social context modulates reward encoding: depending on inequity type, reward encoding was switched on and off rapidly within electrodes, across trials. These results provide direct evidence for explicit encoding of self- and other- rewards in the human OFC, and for rapid and reversible changes in encoding schemes driven by socially relevant contexts.

## Introduction

We often cannot help but compare our own outcomes to those of others: did our sibling get a bigger piece of cake, our friend a better price on her car, our colleague a higher bonus? Comparing rewards is a part of social life, and individuals’ satisfaction with their own outcomes often varies as a function of the outcomes obtained by comparable others (Suls et al., 2002). Inequity aversion, or the preference for a fair, symmetric reward distribution among individuals, is observed widely in human society (Fehr & Schmidt, 1999) - with human children as young as three years old reacting to unequal distributions of rewards (Lobue et al., 2009) - as well as in other primates (Brosnan & de Waal, 2003; Proctor et al., 2013). Navigating decisions involving inequity relies on the interplay of reward valuation and social cognition. A wealth of functional neuroimaging and lesion evidence has pointed to the involvement of the human orbitofrontal cortex (OFC) in both value-based decision-making and social cognition. Here we addressed how the OFC links both processes by conducting intracranial recordings in neurosurgical patients while they performed a social decision-making task.

Historically, lesion studies point to a critical role for the OFC in social functioning. OFC lesions in humans lead to impairments in social judgments and social behaviors, including disinhibition, taking socially inappropriate actions, and misinterpreting others’ moods (Hornak et al., 2003; Perry et al., 2016; Willis et al., 2010, Rolls et al., 1994, Szczepanski & Knight, 2014). Additionally, patients with orbitofrontal damage have difficulty with theory of mind (Stone et al., 1998) and fail to respond to socially-charged stimuli despite retaining normal autonomic responses to other charged stimuli, such as loud noises (Damasio et al., 1990). Finally, human neuroimaging studies show OFC activation during social tasks (Lin et al., 2012; Rilling et al., 2002; Seo et al., 2013; Zaki et al., 2011,).

The role of the OFC in individual value-based decision-making is well established with converging evidence from lesion, neuroimaging, and electrophysiological studies (Wallis, 2007). The OFC encodes a wide range of valuation-related variables such as probability, reward magnitude, prior expectations, and regret (Camille et al., 2004; Domenech et al., 2020; Manssuer et al., 2022; Padoa-Schioppa & Assad, 2006; Saez et al., 2018). Notably, the medial OFC (mOFC; vmPFC) computes the subjective value (Chib et al., 2009; Hunt et al., 2012; Kable & Glimcher, 2007; Lopez-Persem et al., 2020; Plassmann et al., 2007; Strait et al., 2014). Such studies led to the proposal that the OFC is computing a common neural-reward currency by integrating multiple parameters, a useful mechanism to make decisions between options with different attributes (Chib et al., 2009; Levy & Glimcher, 2012).

Despite the numerous studies pointing to the role of the human OFC in both social functioning and value-based decision making, and the clinical relevance of understanding the neural mechanisms of social valuation, little information is available on the fine-scale neuronal processing of social information and its influence on valuation mechanisms. The single-neuron primate literature provides some answers. In one study, macaque monkeys worked to collect rewards either for themselves only or for themselves and a monkey partner. Value coding neurons (e.g., neurons increasing their firing rate with the magnitude of reward) were found to decrease their discharge rate when the monkeys were working to obtain rewards for both themselves and a partner monkey, vs. when they worked to obtain a reward for themselves only. This modulation of the OFC is in line with the behavioral finding that monkeys preferred working in the non-social context. Furthermore, neuronal activity was found to track the identity of the partner monkey (Azzi et al., 2012). The primate orbitofrontal cortex thus contains key neuronal mechanisms for the evaluation of social information. In another study, however, the OFC was found to encode only the monkey’s own reward, independently of the social context (Chang et al., 2013). In humans, iEEG provides the rare opportunity to examine neural mechanisms at a “mesoscale” level of analysis lying between the extensive anatomical coverage of fMRI and the superb temporal resolution of single-unit recordings (Voytek et al., 2015). Here we leverage iEEG to investigate how the OFC makes monetary decisions in different inequity-defined social contexts.

We collected intracranial recordings from 10 neurosurgical patients with electrodes implanted in the OFC while they played a repeated trial Dictator game. In dictator games, a single player (the “dictator”) decides how to split different pots of money between themselves and a social counterpart. Dictator games have been widely used in behavioral and neural research to study social decision-making (Forsythe et al., 1994; Gao et al., 2018). Here, we take advantage of two important features of the task. First, its non-strategic nature makes it easy to understand by the participant and ensures that when the game is repeated, the choices are independent since one doesn’t need to anticipate the social counterpart’s behavior. Second, the simplicity of the task facilitates varying important features of the choice faced by a decision-maker, such as the set of possible payoffs (Krupka & Weber, 2013).

In our task, on each trial, patients had to choose between two allocations of money for themselves and an anonymous other: an equitable option that appeared on all trials ($10 for themselves, and $10 for the other), and a second inequitable option which varied from trial to trial. Depending on the values of the inequitable option, our patients encountered two types of social contexts depending on the type of inequity: *advantageous inequity*, in trials in which the inequitable option presented a higher payoff for the patient than for the other player, and *disadvantageous inequity*, in trials in which the inequitable option presented a lower payoff for the patient than for the other player. In addition to manipulating the type of inequity patients encountered, we also manipulated several reward-related variables in the task by varying the payoffs in the inequitable option from trial to trial.

Given the evidence that the OFC is involved in social functioning and in reward valuation, we hypothesized that the OFC 1) it should encode not only self-related rewards, but also rewards pertaining to the social counterpart; 2) it should be sensitive to the type of inequity present in the trial (advantageous vs. disadvantageous); and 3) the type of inequity should modulate the encoding of self and other-related rewards.

## Results

### Social Decision-Making Behavior in Neurosurgical Patients

We recorded intracranial signals from 10 adult patients (5 female, mean age = 35.6, SD = 10.45, 8 right-handed, 1 ambidextral) who were undergoing intraoperative neurosurgical treatment for refractory epilepsy. As electrode placement and treatment decisions are made solely by the clinical team, the number and specific location of electrodes varied across patients. We recorded from a total of X electrodes of which 143 (136 bipolar pairs) were included in the final dataset as artifact-free, OFC electrodes (for details, See Methods).

Testing was conducted for a single 15-20 minute session and the patient’s pain, medications, and rest was carefully monitored to ensure testing only commenced when the patient was fully alert, capable, and cooperative. On each trial, patients chose between two allocations of money for themselves and another anonymous player (Saez et al., 2015). Figure 1b illustrates the experimental paradigm. Trials started with a fixation cross (t = 0), followed by the game presentation screen (t = 750ms). At that time, patients were given up to 5 seconds to choose between two allocations: a fixed, *equitable* option that did not vary across trials ($10 for themselves; $10 for the other player), and an *inequitable* option that varied across trials. Inequitable trials were w either advantageous (i.e., with a higher payoff for the patient than for the other player, for example, $12 for themselves; $8 for the other player) or disadvantageous (i.e., with a lower payoff for the patient than for the other player, for example, $8 for themselves; $14 for the other player).

**Figure 1:**
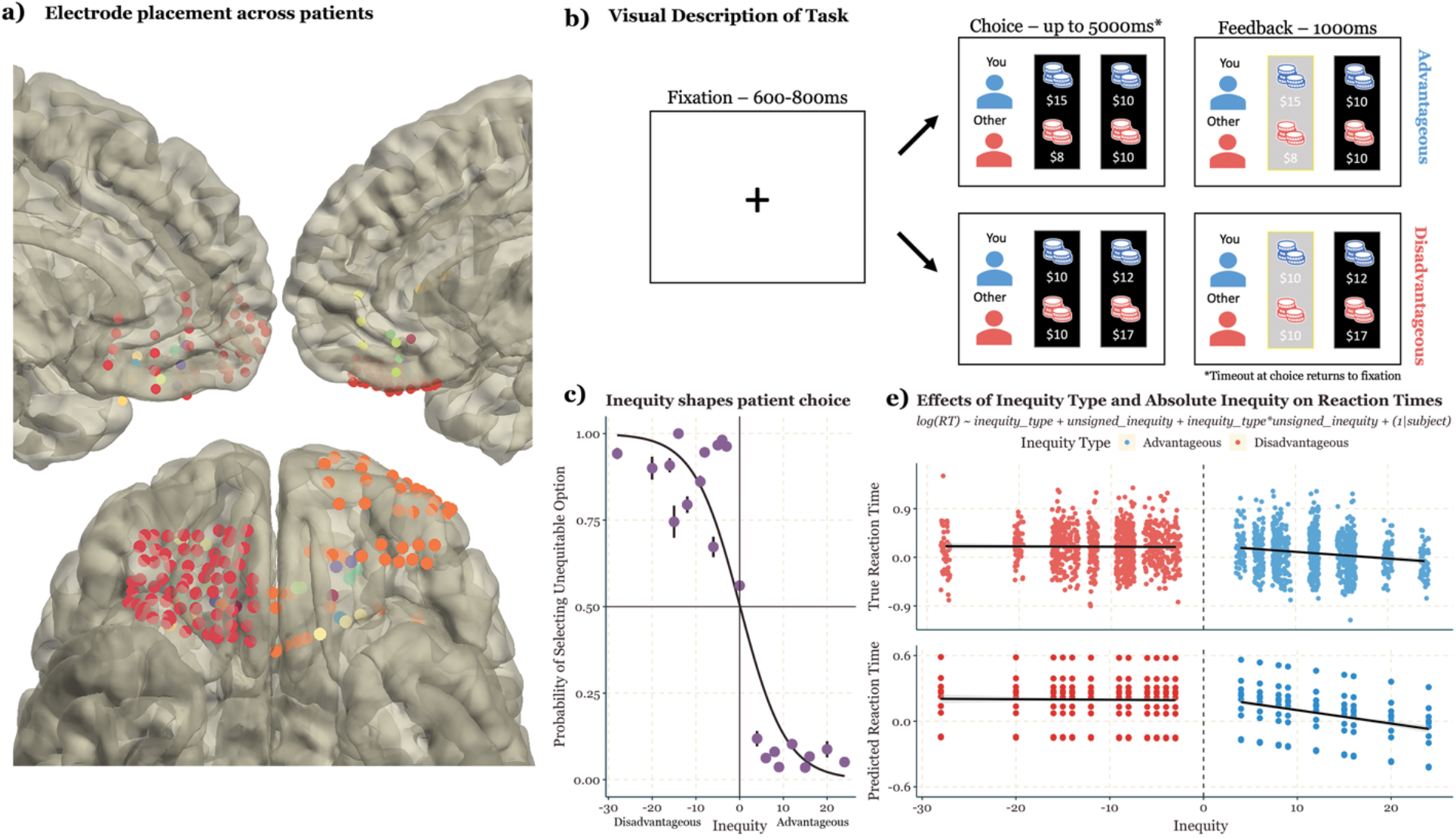
Experimental Approach. **a)** Anatomical reconstruction showing placement of all 143 electrodes in OFC across all 10 patients. Each color corresponds to a patient. **b)** Patients (n = 10) chose between two allocations of money for themselves and an anonymous other player: an equitable option that appeared on all trials ($10 for themselves, and $10 for the other player), and a second inequitable option which varied from trial to trial. Depending on the values of the inequitable option, patients encountered two types of inequities: advantageous inequity, in trials in which the inequitable option presented a higher payoff for the patient than for the other player (see example on the upper row), and disadvantageous inequity, in trials in which the inequitable option presented a lower payoff for the patient than for the other player (see example in the lower row). **c)** Patients’ choices were affected by the signed inequity in the inequitable option (p < 0.0001; random effects logit analysis; error bars = standard error of the mean). **d)** Log-scaled reaction times were predicted by both inequity type and unsigned inequity. Specifically, higher inequity in advantageous trials predicted faster reaction times, while there was no effect of inequity in disadvantageous trials. The top plot shows the true log scaled reaction times on the y axis and signed inequity along the x axis. The bottom plot shows the log scaled reaction times as predicted by our model (Inequity Type: β = -0.035 ± 0.029, t = -1.22 df = 1902; inequity: β = -0.013 ± 0.002, t = -7.40, df = 1902, interaction: β = 0.013 ± 0.002, t = 5.77, df = 1902; random effects analysis).

As expected, all ten patients’ behavior on the task was driven by both the amount offered for themselves in the inequitable option (termed Self-Offer), as well as the signed inequity, (the difference between their offer and the other player’s offer within the inequitable option; p < 0.0001, β = 0.2 ± 0.008, z = 24.77, df = 2159, logistic linear mixed-effect model; Figure 1C). Additionally, patients were particularly sensitive to disadvantageous inequity, even forgoing higher Self-Offer when disadvantageous inequity was high (i.e., $12 Self, $18 Other; 89% +-14% selected the equitable option on disadvantageous trials across patients). While there are other behavioral strategies reported in the literature (i.e., inequity minimizing, self-payoff maximizing), all ten patients in this study were consistent in their choices to minimize disadvantageous inequity (Andreoni & Miller, 2002).

In addition to their choices, the patients’ reaction times were associated with the absolute amount of inequity, demonstrating they were tracking the changing values in the inequitable option. Using a linear mixed-effects model, with fixed effects of Inequity Type, Unsigned Inequity (the absolute value of the difference between their offer and the other player’s offer within the inequitable option) and their interaction and a random effect of patient, we found that log-scaled reaction times were predicted by both Inequity Type and Unsigned Inequity. Specifically, higher inequity in advantageous trials (i.e. $16 Self, $8 Other) predicted faster reaction times, while there was no effect of inequity in disadvantageous trials. (Inequity Type: β = -0.035 ± 0.029, t = -1.22 df = 1902; Unsigned Inequity : β = -0.013 ± 0.002, t = -7.40, df = 1902, Interaction: β = 0.013 ± 0.002, t = 5.77, df = 1902; Figure 1D).

### Social decision variables encoded in OFC via HFA

Given the putative role of the OFC in decision-making, we expected the OFC to encode information about rewards for oneself. Given the role of OFC in social processing, we hypothesized that OFC should encode information about rewards for the social counterpart as well. Following the approach laid out in Saez et al. 2018, we tested our hypothesis that these reward-related, social decision-making variables were encoded via the high-frequency activity in the OFC. Briefly, our regression approach extracted temporal intervals where a variable of interest (Self-Offer, Other-Offer, etc) correlated with the HFA above a permuted null distribution (See Online Methods for full details). As predicted, we found that electrodes encoded Self-Offer across all three epochs of the task (19% of OFC electrodes across task; by epoch, presentation: 5%, pre-choice: 4%, post-choice: 10%; see Figure 2.a, 2.b). Furthermore, as hypothesized, we found that Other-Offer was similarly encoded across many electrodes (30% of OFC electrodes across task; by epoch, presentation: 5%, pre-choice: 10%, post-choice: 15%; see Figure 2A, 2B), though rarely within the same electrode as Self-Offer (See Figure 2B).

**Figure 2:**
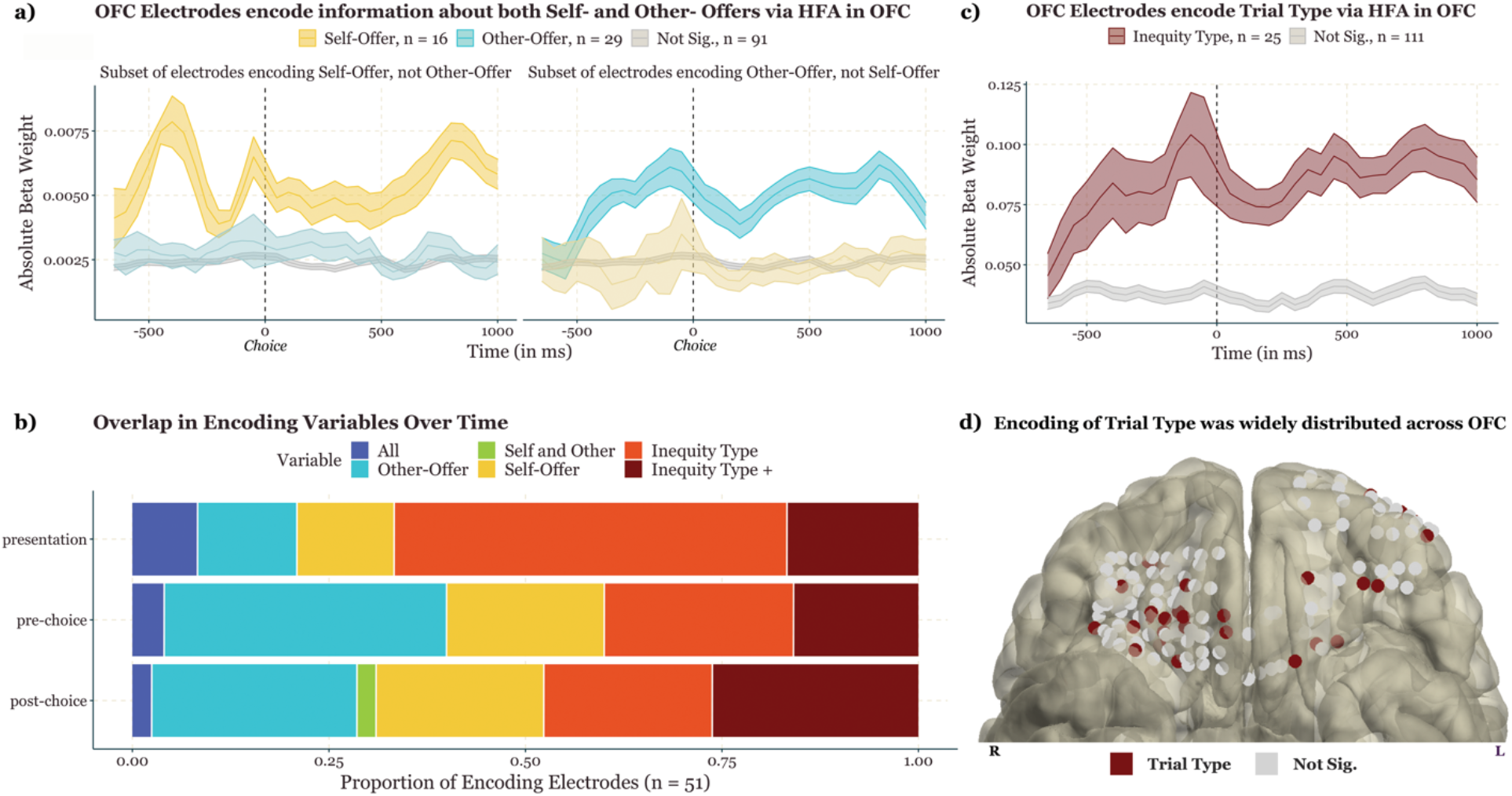
Social decision variables encoded via OFC via HFA. **a)** HFA in OFC electrodes encoded reward-related variables pertaining to both the self and the other player. In the left and right plots, the y axis represents the absolute value of the average beta weights from the regression predicting HFA as a function of Self-Offer and Other-Offer, respectively. Left plot shows all electrodes significantly encoding the Self-Offer (n= 16, yellow), the average beta weight for those same electrodes when Other-Offer was used as a predictor (not significant, blue), and the average beta weights for all electrodes that did not significantly encode Self-Offer nor Other-Offer (n = 91, gray). The right plot similarly shows the average absolute value of beta weights for electrodes significantly encoding Other-Offer (n = 29, blue), those same electrodes when Self-Offer was used as a predictor (not significant, yellow), and the average beta weights for all electrodes that did not significantly encode Other-Offer nor Self-Offer (n = 91, gray). The shaded portions for each line show the standard error of the mean. The dashed vertical line (t=0) indicates the time patients made a choice between the two options. **b)** Social decision-making variables were broadly encoded in OFC electrodes across the three task epochs (presentation, pre-choice and post-choice; see Methods). X-axis shows the proportion of encoding electrodes (n = 51, 37.5% of OFC electrodes) that encoded at least one variable. Self-Offer and Other-Offer were encoded robustly across all three epochs, but rarely did the same electrode encode both those variables during the same epoch (3.6% of encoding electrodes, including those that also encoded Inequity Type). Inequity Type was also robustly encoded across all three epochs and was frequently encoded along with either Self-Offer or Other-Offer, denoted ‘Inequity Type +’. Finally, 2.9% of encoding electrodes encoded all three variables within a single epoch. **c)** Inequity Type was encoded via HFA in the OFC. The figure plots the average absolute value of the beta weights across all electrodes that significantly encoded Inequity Type (n = 25, red) and across electrodes that did not encode Inequity Type (n = 111, gray). The dashed vertical line (t=0) indicates the time patients made a choice between the two options. The shaded portions for each line show the standard error of the mean. **d)** Anatomical localization of electrodes encoding Inequity Type (dark purple dots; gray dots represent non-encoding electrodes). See SOM for localization for other variables.

Next, since our study, in line with previous work, demonstrates that inequity type moderates patients’ behavior, we assessed if Inequity Type (e.g., whether the current trial was an Advantageous inequity or Disadvantageous inequity one) was also encoded in the OFC via HFA. Importantly, as a binary variable, Inequity Type is less related to the amount of reward or value associated with the trial, but instead captures the social context of a given trial. Using a similar approach as with Self- and Other-Offer, we found many electrodes significantly encoded Inequity Type across all three task epochs (37% of ofc electrodes across task; by epoch, presentation: 13%, pre-choice: 8%, post-choice: 15%; see Figure 2B, 2C). Additionally, we found that a single electrode frequently encoded Inequity Type along with either Self-Offer or Other-Offer (see Figure 2B).

### State-dependent Inequity Encoding results

Based on the predominance of the Inequity Type effect we hypothesized that Inequity Type would moderate how Self-Offer and Other-Offer were encoded in the HFA. In a secondary analysis, we thus looked at the encoding of different variables separately in advantageous and disadvantageous trials. There were three different possibilities from these analyses. First, we could see comparable encoding strengths in each inequity type, indicating inequity type does not modulate Self-Offer and Other-Offer encoding. Second, the encoding strength could be diminished in one inequity type compared to another. For example, Self-Offer might be encoded less strongly in disadvantageous trials compared to advantageous trials, perhaps reflecting a devaluation of the offer in the face of large disadvantageous inequity as reported in the behavioral literature (Suls et al., 2002). Third, specific electrodes might encode reward variables only in (dis)advantageous trials, and “switch off” when the patient is experiencing the other type of inequity, which we call state-dependent inequity encoding. We found that many of the electrodes in this sample used this third encoding scheme, providing clear evidence against the first option, that there is no modulation of inequity encoding on reward variable encoding. We were not able to evaluate the evidence for the second encoding type, due to the statistical limitations of testing for interaction terms within a permutation framework (Edgington, 1995; Jung et al., 2007; Ter Braak, 1993; Still & White, 1981)(see methods for details). Importantly, an effect for the second encoding possibility could not explain the state-dependent inequity encoding results.

To test for the third encoding type in the full sample, we ran regressions predicting the HFA using each predictor in advantageous and disadvantageous trials separately. We used the absolute value of the summed difference in the R2 lines as a permutation statistic. This statistic is small whenever either the regressor poorly predicts the HFA in both trials and when the regressor well predicts the HFA in both trials but is large when the R2 values are high in only one Inequity Type. We found that many electrodes encoded Self-Offer (37% of ofc electrodes across task; by epoch, presentation: 10%, pre-choice: 13%, post-choice: 14%) and Other-Offer (45% of ofc electrodes across task; by epoch, presentation: 13%, pre-choice: 14%, post-choice: 18%) in a state-dependent manner, where encoding would “turn on” and “turn off” depending on the Inequity Type. Additionally, we found that an electrode’s preferred Inequity Type was consistent within the task. In other words, if an electrode began encoding a variable in only advantageous trials, it would continue to encode only in the advantageous trials for the remaining epochs, even if the predictor it encoded changed (see Figure 3B). Only one electrode switched from encoding in the advantageous trials to the disadvantageous trials over the course of a trial.

**Figure 3:**
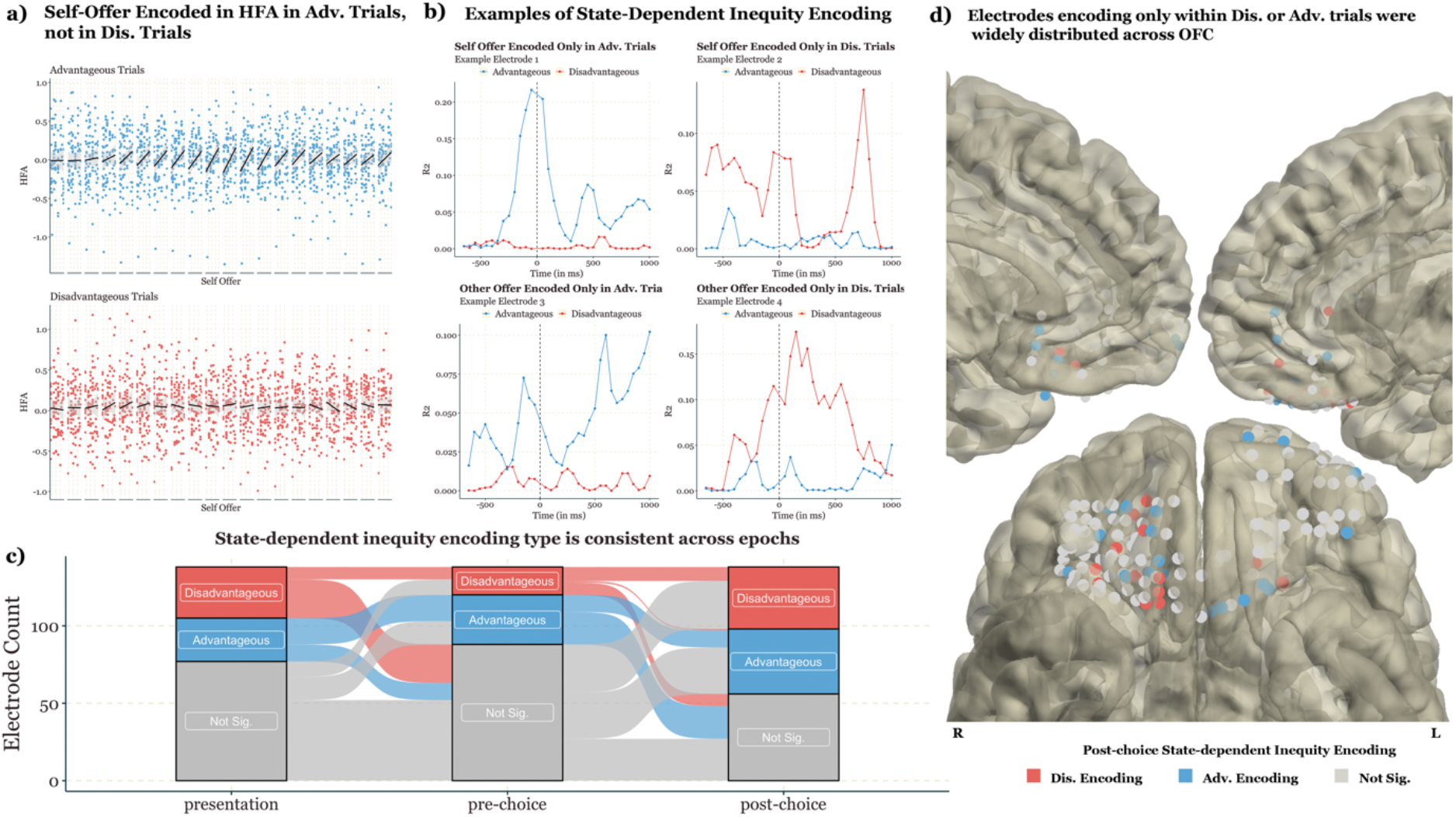
State-Dependent Inequity Encoding. **a)** Example of an electrode encoding Self-Offer in Advantageous trials, but not in Disadvantageous trials. The top plot shows the individual regressions across each time bin (50ms) with the X axis showing the Self-Offer values presented across trials and the Y axis showing the HFA baselined to 200ms before trial onset and each dot represents an advantageous trial. The bottom plot is as above, but for the disadvantageous trials. While Self-Offer was significantly encoded in advantageous trials, no such relationship was detected for disadvantageous trials. **b)** Time course of encoding in example electrodes. Each subplot shows the R2 from the individual regression for both advantageous (blue lines) and disadvantageous (red lines) trials for example electrodes. The top and bottom plots show example electrodes that encoded Self-Offer and Other-Offer, respectively. The left and right plots show encoding in advantageous and disadvantageous trials, respectively. **c)** Inequity encoding type across epochs. Y-axis represents the percentage of electrodes that encoded a particular reward-related variable, and the facets show how this percentage changed across the three task epochs. Electrodes that encoded task variables (Self-Offer, Other-Offer, Max, and Min) in advantageous trials tended to not encode task variables in disadvantageous trials in different epochs. **d)** Anatomical localization of state-dependent inequity encoding electrodes in the post-choice epoch (blue/red/gray dots represent electrodes encoding only in advantageous/disadvantageous/none conditions).

Importantly, we evaluated alternative explanations that might be driving this effect. First, within these inequity-dependent encoding electrodes, we examined if one inequity type was noisier than the other, which could obscure the encoding in one of the inequity types. To test this, we extracted the residuals from each time bin in a given epoch and performed a t-test between the two inequity types. If the confidence interval of the difference in means did not include 0, we excluded this electrode’s epoch from the inequity-dependent analyses. We found that differences in noise between the two conditions were rare, occurring in only 1.5% of electrode-epoch pairs, suggesting that the state-dependent inequity encoding is not driven by differences in noise in the underlying neural signal. In addition, to minimize the potential impact of such an effect, we removed these electrodes from subsequent inequity-dependent analyses.

It is possible that these state-dependent inequity encodings could be driven by a “unified” encoding of a different predictor. By “unified” we mean that the predictor was encoded across both inequity types similarly. For example, Self-Offer is identical to Maximum Value in advantageous trials, meaning that electrodes that seem to be encoding “Self-Offer” in only advantageous trials might instead be encoding Maximum Value across all trials. To address this possibility, we tested each state-dependent electrode that encoded either Self-Offer in Advantageous trials or Other-Offer if Disadvantageous trials to see if those electrodes also significantly encoded Max Value across both inequity types. In total, there were five state-dependent encoding electrodes whose behavior could also be explained by the unified encoding of Maximum Value. These electrodes were excluded from the remaining state-dependent analyses. The same process was conducted to see if Minimum Value unified encoding could explain Self-Offer encoding in Disadvantageous trials and Other-Offer encoding in only Advantageous trials. None of these electrodes’ responses were significantly predicted by Minimum value encoding across both inequity types.

### Anatomical Distribution of Social Reward Encoding

We assessed whether any anatomical organization within the OFC was linked to all Self-Offer, Other-Offer, and Inequity Type encoding. We tested this by predicting the test statistic from our electrode-specific linear models (see Online Methods for details) from each of the three averaged MNI coordinates of the bipolar-referenced electrodes. We found a modest effect of electrode location within our state-dependent analyses. Specifically, the Y coordinate (anterior/posterior axis) predicted the magnitude of the state-encoding effect across all the predictors (p = 0.03, β = 0.001 ± 0.0005, t = 2.161, df = 1603, linear mixed-effect model). Thus, there was a higher likelihood of state-dependent inequity encoding the more anterior the electrode was placed. However, this result does not survive multiple comparison correction. Furthermore, there were no effects of anatomical localizations along the X (lateral/medial) axis) or Z (superior/inferior axis) axes. Overall, this is consistent with the findings that HFA encoding of different reward-related variables is distributed across the OFC and vmPFC (Saez et al., 2018).

## Discussion

The role of the orbitofrontal cortex in value-based choice is well-established, but its role in human social decision-making is less clear. We report to our knowledge the first findings from human intracranial study on social decision-making and inequity processing. We recorded iEEG signals from patients playing a repeated dictator game in which they chose between two allocations of money for themselves and another anonymous player: one fair, equitable option, and one inequitable option – either advantageous or disadvantageous. Our analyses focus on High-Frequency Activity, an index of local cortical computation (Buzsáki et al., 2012; Leszczynski et al., 2020) that captures value-based computations in non-social tasks (Domenech et al., 2020; Saez 2018). We provide novel evidence that OFC HFA encodes social reward-related variables and provide further evidence for state-dependent inequity encoding, delineating a new mechanism by which the OFC adapts rapidly to changing social contexts.

### Behavior

Some studies report that individuals dislike advantageous inequity and are willing to augment others’ incomes at their own personal cost (Dawes et al., 2007; Yu et al., 2014), although significant inter-subject variability exists (Saez et al., 2015). Our patients’ choices suggest that they attempted to minimize disadvantageous inequity even at the expense of their own payoff but didn’t care about minimizing advantageous inequity. For example, if offered $12 for themselves and $16 for their social counterpart, they would frequently select the equitable option ($10 for both themselves and their counterpart), despite the smaller payout for themselves. Importantly, reaction times analysis revealed that patients were also responding to the amount of advantageous inequity: they had faster choices the bigger the advantageous inequity, suggesting they experienced less conflict (Evans et al., 2015). The homogeneity of choices in our sample could be due to its size (N=10), which is at the high end of the range for iEEG studies (Domenech et al., 2020), but smaller than behavioral and neuroimaging studies reporting advantageous inequity aversion. Importantly, these considerations mean that the results we report may be representative of participants whose choices are swayed by inequity type but might be less representative of participants who disregard inequity to either maximize their own payout or maximize the payout of the whole (Saez et al., 2015).

### The OFC encodes social variables in addition to self-related variables

We found that human OFC, in addition to encoding Self-Offer, also encodes variables relevant to their social counterpart: 30% of the OFC electrodes tracked the value of Other-Offer. This is the first evidence for encoding of other-reward in the human OFC extending the findings that high-frequency activity in human OFC encodes self-related reward variables in a non-social decision-making task (Saez et al., 2018). These results also compliment the non-human primate literature. Several studies have reported that OFC neurons only encode rewards delivered to oneself, and not to another monkey, while neurons in other regions such as the anterior cingulate gyrus encoded both (Azzi et al., 2012; Chang et al., 2013). Here we show human OFC encodes both self and other rewards.

### Inequity type is encoded in the OFC

We found that the OFC was sensitive to the type of inequity (advantageous vs. disadvantageous) presented in the trial: 36% of our sample’s OFC electrodes encoded inequity type, an effect observed in 9 of our 10 patients. It is worth emphasizing that inequity type was not explicitly mentioned in the game instructions nor signaled to our patients in the trial display (i.e., with a different background color). The predominance of the effect suggests that when presented with the inequitable option, patients rapidly engaged in a comparative process of the Self- and Other-Offers, confirming the importance of fairness considerations in valuation and decision-making. Furthermore, this variable represents an example of social context encoding in the OFC, one that was relevant to our patients’ behavior, but does not explicitly include reward values.

### State-dependent inequity encoding of reward variables

The prominent Inequity Type effect and our behavioral findings prompted investigation of whether inequity type might modulate the encoding of other reward-related variables. We examined the encoding of different variables separately in advantageous and disadvantageous trials and observed a novel type of encoding in the OFC – which we refer to as “state-dependent inequity encoding”. We found that substantial numbers of OFC electrodes encoded at least one variable in an inequity-dependent way: 31% of the electrodes showed advantageous only encoding - encoding at least one variable only in advantageous trials - and 35% showed disadvantageous only encoding -encoding at least one variable only in disadvantageous trials. The state-dependent inequity encoding was found in each one of our 10 patients. This state-dependent inequity encoding shows that the OFC flexibly and rapidly adapts to changes in context, as in our study, the inequity context changed from trial-to-trial.

State-dependent inequity encoding provides a new mechanism by which social context impacts reward encoding differing from the modulation effect of social context found in past studies. In the Azzi et al.’s study, where monkeys worked in different social contexts (to collect rewards for themselves only, or for themselves and a partner monkey), neurons encoding reward value were found to have a reduced activity when the monkeys were offered to work for a joint reward, mirroring the behavioral observation that they were less motivated to work in the social condition (Azzi et al., 2012). However, in our human iEEG study inequity type did not simply modulate the encoding of a specific variable (for example by increasing the relationship between HFA and Self-Offer in advantageous vs. disadvantageous trials), but rather turned on and off this encoding in distinct populations of OFC electrodes. Importantly, the two types of modulation (state-dependent and decrease of encoding) are not mutually exclusive and could coexist in different OFC electrodes.

Our findings are consistent with the recent idea that the OFC contains a “cognitive map” of task space in which the current state of the task is represented (Wikenheiser & Schoenbaum, 2016; Wilson et al., 2014). This idea was initially supported by a study showing that patterns of fMRI activity in human OFC contain information about subjects’ current location in a cognitive map of a task composed of 16 different hidden states (Schuck et al., 2016; for similar results in rodents’ OFC, see Zhou et al., 2019). The two states of our Dictator task (advantageous and disadvantageous inequities) were clearly encoded, suggesting that the notion of “task states” can be expanded to encompass social contexts as well. This is reminiscent of findings showing that the hippocampus encodes ‘‘social space’’ (i.e., power and affiliation) in the form of a cognitive map (Schafer & Schiller, 2018), and in line with the evidence that the OFC and the hippocampus’ cognitive mapping are similar (Wikenheiser & Schoenbaum, 2016). Importantly, our findings on context-dependent encoding provide a neural mechanism by which the current state of a task influences further computations.

## Conclusion

We found that the OFC is sensitive to social context. In addition to encoding self-related variables (such as the offer for self), OFC also encodes the offer for the social counterpart, as well as the type of inequity (advantageous vs. disadvantageous). We also found a new mechanism by which social context modulates the encoding of these variables. Depending on inequity type, reward encoding was switched on and off in distinct orbitofrontal electrodes. This context-dependent encoding provides a potential mechanism by which the OFC constructs a “cognitive map of task space”, and flexibly and rapidly adapts valuation computations to different task states. Social context-dependent encoding may not be specific to inequity. In fact, the value of decisions we make each day is dependent on the social context we are in– having a glass of wine at dinner among friends is frequently deemed an acceptable choice, while doing so in the middle of a lecture is not. Future research is needed to uncover how other social contexts, such as the social distance between individuals (Fareri et al., 2012; Schreuders et al., 2018) or the cooperative vs. competitive nature of the interaction (Bault et al., 2011), are encoded.

## Methods

### Patients

We recorded intracranial signals from 10 (5 female) adult patients (mean age = 36.6 years, SD = 10.45age range = 25–58 years) who were undergoing intraoperative neurosurgical treatment for refractory epilepsy. Each patient was implanted with subdural grids, strips and/or depth electrodes to localize the seizure onset zone for subsequent surgical resection. We selected patients with electrodes implanted in the OFC/vmPFC (range, average, overall). The cohort consisted of X unilateral cases (Y left, W right) and Z bilateral cases. The position of these electrodes was dictated solely by the patient’s clinical needs. Patient recordings took place at four hospitals: the University of California, San Francisco (UCSF, N=1) Hospital, the Stanford School of Medicine (N=1), the University of California Irvine Medical Center (UCI, N=7) and the University of California, Davis (Davis, N=1). All patients provided written informed consent as part of the research protocol approved by each hospital’s Institutional Review Board and by the University of California, Berkeley. Patients were tested when they were fully alert and cooperative.

### Behavioral task

We investigated social preferences using a modified Dictator task in which patients chose between two allocations of money for themselves and another anonymous player (Saez et al., 2015). Figure 1b illustrates the experimental paradigm. Trials started with a fixation cross (t = 0), followed by the game presentation screen (t = 750ms). At that time, patients were given up to 5 seconds to choose between two allocations: a fixed, EQUITABLE option that did not vary across trials ($10 for themselves; $10 for the other player), and an INEQUITABLE option, that varied across trials. The inequitable was either advantageous (i.e., with a higher payoff for the patient than for the other player, for example, $12 for themselves; $8 for the other player) or disadvantageous (i.e., with a lower payoff for the patient than for the other player, for example, $8 for themselves; $14 for the other player). In addition, as a control we included catch trials (n=??) in which the second allocation was also equitable and dominated the fixed $10/$10 option (for example $14 for themselves; $14 for the other player). Once a choice was made (using the ?? or ??? keys on the keyboard), the chosen allocation was highlighted for 1000ms, after which a new trial started. If the patient did not choose within the allotted time limit, a timeout occurred and the game moved on to the following trial. Timeouts were infrequent (4% +-1% of trials across patients) and were excluded from the analysis. The location of the equitable and non-equitable allocations (left/right) was randomized across trials.

Patients were instructed that there was no wrong or right way to play the game, and that they should play according to their own preferences. Due to IRB limitations, we were unable to pay patients according to their decisions in the game, but we asked them to make decisions as if they were playing with real money. Patients completed a training session prior to the game in which they played at least X rounds under the experimenter’s supervision until they felt confident that they understood the task. The game itself was composed of 228 trials. A full experimental run typically lasted approximately 20 minutes. Stimulus presentation was operated by pygame.

### Data Acquisition

Electrophysiological data were recorded using Tucker-Davis Technologies (Stanford, and UCSF), Nihon-Kohden (UCI) or Natus (Davis) systems. Data processing was identical across all sites: channels were amplified ×10000 and analog filtered (0.01-1000 Hz) with > 2kHz digitization rate. The photodiode’s input was recorded in the electrophysiological system as an analog input. This signal was used to synchronize the behavioral and electrophysiological data.

### Data preprocessing

Offline, continuous data were downsampled to 1KHz, low-pass filtered at 200Hz, and notch-filtered at 60Hz and its harmonics up to 300 Hz to remove line noise (Butterworth, 4th order, 2 Hz bandwidth). The data was then demeaned and detrended, before being re-referenced, using a common average reference for grids and strips, and a bipolar reference to an adjacent electrode for depth electrodes. We visually identified and removed channels with poor contact or excessive noise throughout the recording. In addition, each dataset was visually inspected with a neurologist in order to remove electrodes exhibiting epileptic activity and noisy epochs (such as epochs with a spread of epileptic activity from the primary epileptic site). HFA was extracted using a bandpass-Hilbert approach (Bruns, 2004) to extract HFA, 20-Hz-wide sub-bands spanning from 70 to 150 Hz in 5 Hz steps (70 to 90, 75 to 95, … up to 130 to 150 Hz). Finally, we segmented the continuous EEG data into three epochs. Presentation is defined as trial onset until 750ms, pre-choice is defined from 650ms until button press, and post-choice is defined from button press until 1000ms. Each epoch was baselined using the first 200ms preceding trial onset. Data preprocessing was carried out in Matlab using the Fieldtrip Toolbox (Oostenveld et al., 2011). Data analysis was carried out in R using custom scripts.

### Anatomical reconstructions

We used an anatomical data processing pipeline (Stolk et al., 2018) to localize electrodes from a pre-implantation MRI and a post-implantation CT scan. The MRI and CT images were aligned to a common coordinate system and fused with each other using a rigid body transformation. We then compensated for brain shifts caused by the implantation surgery. A hull of the patient brain was generated using the Freesurfer analysis suite. Electrodes were then classified by a neurologist according to the anatomical location within each subject’s anatomical space. Only electrodes confirmed to be in the OFC/vmPFC were included in the analysis. Out of these 144 OFC electrodes, 134 were artifact-free and included in subsequent analyses (range 2-60 per patient, mean 13.4 electrodes). For illustration purposes, we converted patient-space electrodes into Montreal Neurological Institute (MNI) coordinates using volume-based normalization. Fig. 1 shows all OFC/vmPFC used in the analysis.

|  | Variable Name | Qualitative Description | Quantitative Description |
| --- | --- | --- | --- |
| *Primary Variables Of Interest* | Self-Offer | The amount of money offered for oneself in the inequitable offer. *(The amount offered for the self in the equitable option is always \$10, and was not analyzed)* | – |
| | Other-Offer | The amount of money offered for the social counterpart in the inequitable offer. *(The amount offered for the counterpart in the equitable option is always \$10, and was not analyzed)* | – |
|  | Inequity Type | A binary variable denoting whether the trial included an inequitable option where the self could receive a higher payout than the social counterpart OR where the social counterpart could receive a higher payout than oneself | *if*<br>*Self-Offer > Other-Offer,*<br>*Advantageous*<br>*else*<br>*Other-Offer > Self-Offer,*<br>*Disadvantageous* |
| *Secondary Variables of Interest* | Max | The maximum value of the inequitable option | *max(Self-Offer, Other-Offer)* |
|  | Min | The minimum value of the inequitable option | *min(Self-Offer, Other-Offer)* |
|  | Unsigned Inequity | The absolute value of the difference in potential payout within the inequitable option | *abs(Self-Offer - Other-Offer)* |
|  | Signed Inequity | The difference in potential payout within the inequitable option | *Self-Offer - Other-Offer* |

### Social decision variables encoding in the OFC via HFA

To determine if a given electrode was encoding one of our task-related variables (see SOM for table of variables), we employed a regression approach where the dependent variable was defined as the analytic amplitude of the HFA time series extracted via Hilbert transform (Saez et al., 2018). We then divided HFA time series into three event-related epochs to account for inter-trial latencies: presentation (0-750ms after trial onset), pre-choice (0-650ms before choice), and post-choice (0-1000ms after choice). We used a 200-ms baseline to remove any pre-stimulus differences in baseline amplitude and averaged HFA activity using a 200ms rolling window at 50ms increments. To identify task-selective channels, we performed separate linear regressions of average HFA activity on each reward-related regressor of interest.

We used the resulting R^2^ (variance in the neural data that can be explained by the behavioral regressors of interest, % explained variance, %EV) as a metric of the quality of the fit. This approach is insensitive with respect to time of task-related activation and to the direction of encoding (i.e., HFA increases or decreases). We then calculated the encoding window in individual regressors/electrodes by taking the longest stretch of time in which all time points showed significant encoding (p < 0.05). False-positive rate was determined using a permutation strategy, where the test statistic was the sum of the F statistics across the encoding window. For each regressor-HFA regression, we shuffled the relationship between behavioral labels and HFA activity 1,000 times. The resulting distribution was taken as the null for that regressor-electrode combination.

### State-Dependent Inequity Encoding

To test if electrodes encoded reward variables in one Inequity Type but not the other, we took a similar approach used for unified encoding. We ran each predictor-electrode separately for advantageous-type trials and disadvantageous-type trials. Advantageous-type trials were defined as trials where within the varying option, the Self-Offer was higher than the Other-Offer. Disadvantageous trials were defined conversely, where the Other-Offer was higher than the Self-Offer in the variable option. As above, we took a permutation approach. For each variable-electrode pair, we shuffled the relationship between behavioral labels and HFA activity 1,000 times. To simultaneously capture 1) encoding of a reward-related variable in one Inequity Type and 2) a *lack* of evidence for encoding of a reward-related variable in the corresponding Inequity Type, we defined the test statistic for the permutation test as the absolute difference in R2 values between the two regressions, across the entire epoch. Note that unlike in the unified encoding analyses, this method does not filter to only include significant stretches of encoding. This allows us to avoid testing separately for encoding in the advantageous trials and then again in the disadvantageous trials but doesn’t allow us to capture specific time windows of encoding within an epoch.

We then performed a second permutation test on the electrodes that survived our first test of state-dependent encoding. The first test calculates the chance one would see a given reward-related variable encoded in exactly one Inequity Type by chance. In the second permutation, we test the likelihood of seeing encoding of a given reward-related variable in only one-half of the trials by shuffling the Inequity Type labels. The test statistic was calculated as above, resulting in a second null distribution for each regressor-electrode pair. Electrode-regressor pairs were classified as using state-dependent encoding only if the regression was significant (p < 0.05) by both permutation strategies.

### Alternative hypotheses

To confirm that our state-dependent inequity results could not be better explained by differences in noise between the two inequity types, we performed a secondary analysis that compared the residuals between inequity types for each regression across all time bins. Within state-dependent inequity encoding electrodes, we took the residuals from each time bin in each epoch and performed a t-test between the two inequity types. If the confidence interval of the difference in means did not include 0, we excluded this electrode’s epoch from the inequity-dependent analyses. We found that differences in noise between the two conditions were rare, happening in only 1.5% of electrode-epoch pairs, suggesting that the state-dependent inequity encoding is not driven by differences in noise in the underlying neural signal.

Our second alternative hypothesis was that the state-dependent inequity encoding analysis might actually be detecting unified encoding a different variable, whereby “unified” we mean that the predictor was encoded across both inequity types similarly. For example, instead of an electrode encoding Self-Offer only in Advantageous trials, instead that electrode might be encoding Max Value in both Advantageous and Disadvantageous trials. To evaluate this possibility, we took all the electrodes that encoded Self-Offer in a state-dependent manner and tested to see if they also encoded Max value across all trials, using the regression and permutation approach described above. In total, there were five state-dependent encoding electrodes whose behavior could also be explained by the unified encoding of Maximum Value. These electrodes were also excluded from the remaining state-dependent analyses. The same process was conducted to see if Minimum Value unified encoding could explain Self-Offer encoding in Disadvantageous trials and Other-Offer encoding in only Advantageous trials. None of these electrodes’ responses were significantly predicted by Minimum value encoding across both inequity types.

### Anatomical Analyses

To determine if the effects of our above analyses were localized to a specific region of the OFC we used linear mixed-effects modeling. Specifically, we averaged the MNI coordinates of the bipolar-referenced electrodes for each MNI axis. We then tested if the MNI coordinate predicted the test statistic, using a different model for each MNI axis, each with a random effect of subject. See the above methods for the definition of the test statistic. We ran these analyses for both the unified and state-dependent encoding analyses first grouped across all predictors, and then separately for each of our main predictors (Self-Offer, Other-Offer, Inequity Type).

## Limitations

This paper has limitations. First, we explored predictors drawn from the social decision-making fMRI literature to explain electrode activity using a unified encoding scheme. However, there may be other variables that have not yet been considered by the literature. Second, to address concerns of generalizability of our patients to the general population, we undertook extensive efforts to only test patients fully alert and cooperative and removed from analysis electrodes placed over seizure foci or contaminated by artifacts. Additionally, we addressed potential fatigue issues and the strict time limits of our recording sessions by minimizing the cognitive complexity of the task.

There are additional limitations of the specific implementation of this dictator game for studying state-dependent encoding schemes. First, while time with patients is limited, both increasing the number of trials within each inequity type and ensuring that the range of variables across inequity types was equivalent would be an important next step in follow up analyses. Second, while the inequitable option was pseudorandomly presented on the left- and right-hand side, the option that was most attractive to disadvantageous inequity minimizers was more frequently on the left-hand side (69.1% of trials). As our patients mostly employed a strategy to minimize disadvantageous inequity, they mostly selected the left-hand side option (70.4% +-8% of trials on average). This side bias is unlikely to undermine the state-dependent inequity results because 1) our analyses focused exclusively on variables related to the presentation of offers, not on patients’ choice, 2) the inequitable option was presented as frequently on the left as it was on the right, 3) we found evidence of encoding both Self-Offer and Other-Offer in both inequity types– in other words even though the inequitable option was rarely selected on disadvantageous trials, we still find evidence of electrodes encoding this value (see Figure 3.B).

