## Supplemental Figures and Tables for "To share or not to share? iEEG evidence for state-dependent inequity encoding in the human OFC"

| Variable | Epoch | Number of Encoding Electrodes per Epoch | Percentage of Encoding Electrodes per Epoch | Number of Encoding Electrodes Across All Epochs | Percentage of Encoding Electrodes Across All Epochs |
|---|---|---|---|---|---|
| Inequity Type | presentation | 18 | 13.24% | 50 | 36.76% |
| Inequity Type | pre-choice | 11 | 8.09% | 50 | 36.76% |
| Inequity Type | post-choice | 21 | 15.44% | 50 | 36.76% |
| Self-Offer | presentation | 7 | 5.15% | 27 | 19.85% |
| Self-Offer | pre-choice | 6 | 4.41% | 27 | 19.85% |
| Self-Offer | post-choice | 14 | 10.29% | 27 | 19.85% |
| Other-Offer | presentation | 7 | 5.15% | 42 | 30.88% |
| Other-Offer | pre-choice | 14 | 10.29% | 42 | 30.88% |
| Other-Offer | post-choice | 21 | 15.44% | 42 | 30.88% |
| Min | presentation | 6 | 4.41% | 40 | 29.41% |
| Min | pre-choice | 14 | 10.29% | 40 | 29.41% |
| Min | post-choice | 20 | 14.71% | 40 | 29.41% |
| Max | presentation | 9 | 6.62% | 32 | 23.53% |
| Max | pre-choice | 9 | 6.62% | 32 | 23.53% |
| Max | post-choice | 14 | 10.29% | 32 | 23.53% |
| Unsigned Inequity | presentation | 4 | 2.94% | 14 | 10.29% |
| Unsigned Inequity | pre-choice | 2 | 1.47% | 14 | 10.29% |
| Unsigned Inequity | post-choice | 8 | 5.88% | 14 | 10.29% |

**Supplemental Table 1.** Results from the unified encoding analyses.

| Variable | Epoch | Number of Encoding Electrodes per Epoch | Percentage of Encoding Electrodes per Epoch | Number of Encoding Electrodes Across All Epochs | Percentage of Encoding Electrodes Across All Epochs |
|---|---|---|---|---|---|
| Self-Offer | presentation | 14 | 10.29% | 49 | 36.03% |
| Self-Offer | pre-choice | 17 | 12.5% | 49 | 36.03% |
| Self-Offer | post-choice | 18 | 13.24% | 49 | 36.03% |
| Other-Offer | presentation | 16 | 11.76% | 50 | 36.76% |
| Other-Offer | pre-choice | 15 | 11.03% | 50 | 36.76% |
| Other-Offer | post-choice | 19 | 13.97% | 50 | 36.76% |
| Min | presentation | 9 | 6.62% | 35 | 25.74% |
| Min | pre-choice | 6 | 4.41% | 35 | 25.74% |
| Min | post-choice | 20 | 14.71% | 35 | 25.74% |
| Max | presentation | 22 | 16.18% | 59 | 43.38% |
| Max | pre-choice | 12 | 8.82% | 59 | 43.38% |
| Max | post-choice | 25 | 18.38% | 59 | 43.38% |

**Supplemental Table 2.** Results from the state-dependent inequity encoding analyses.

**Encoding of Self-Offer was widely distributed across OFC**

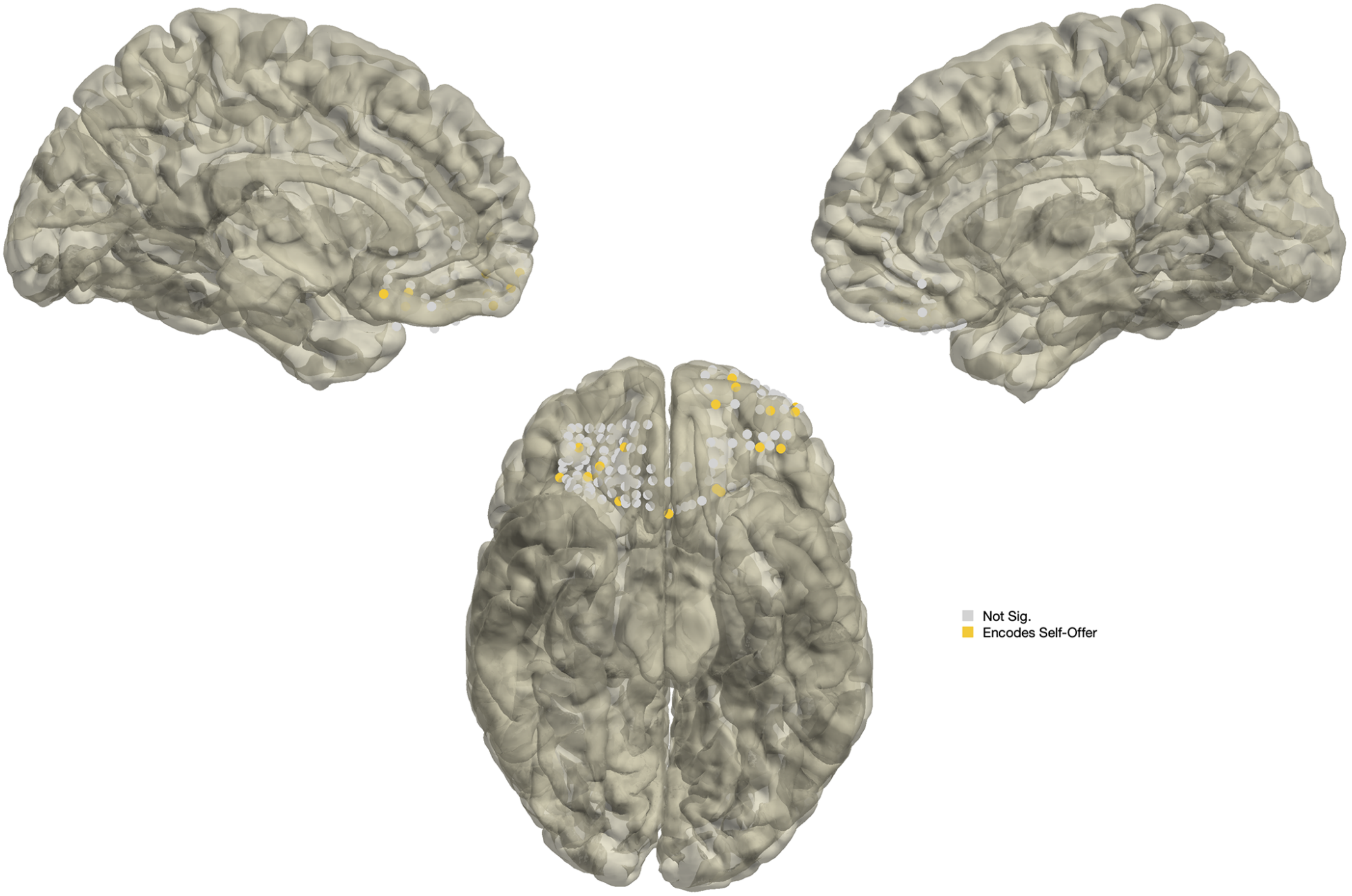


**Supplemental Figure 1.** Anatomical localization of electrodes encoding Self-Offer (yellow dots; gray dots represent non-encoding electrodes).

**Encoding of Other-Offer was widely distributed across OFC**

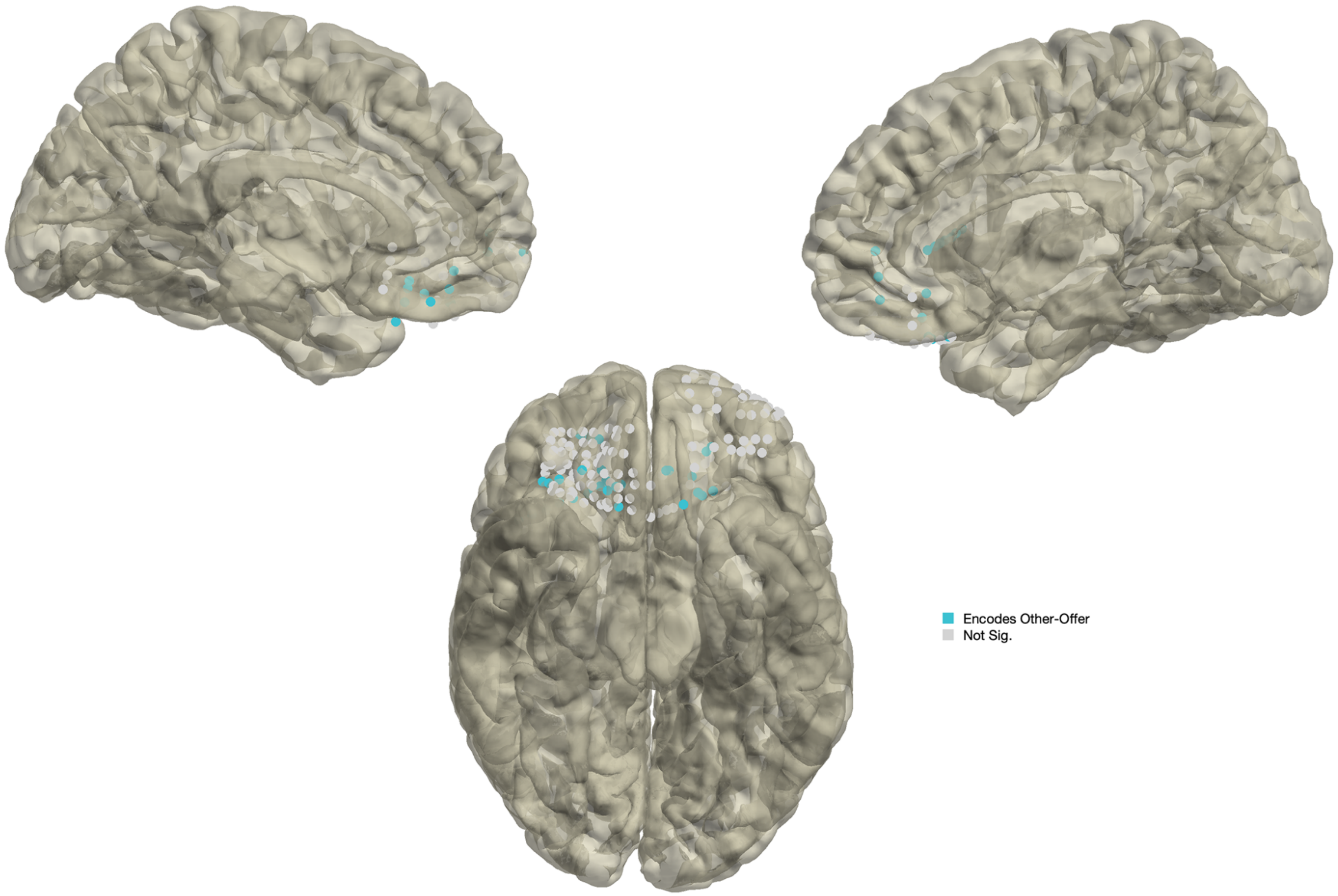


**Supplemental Figure 2.** Anatomical localization of electrodes encoding Other-Offer (blue dots; gray dots represent non-encoding electrodes).

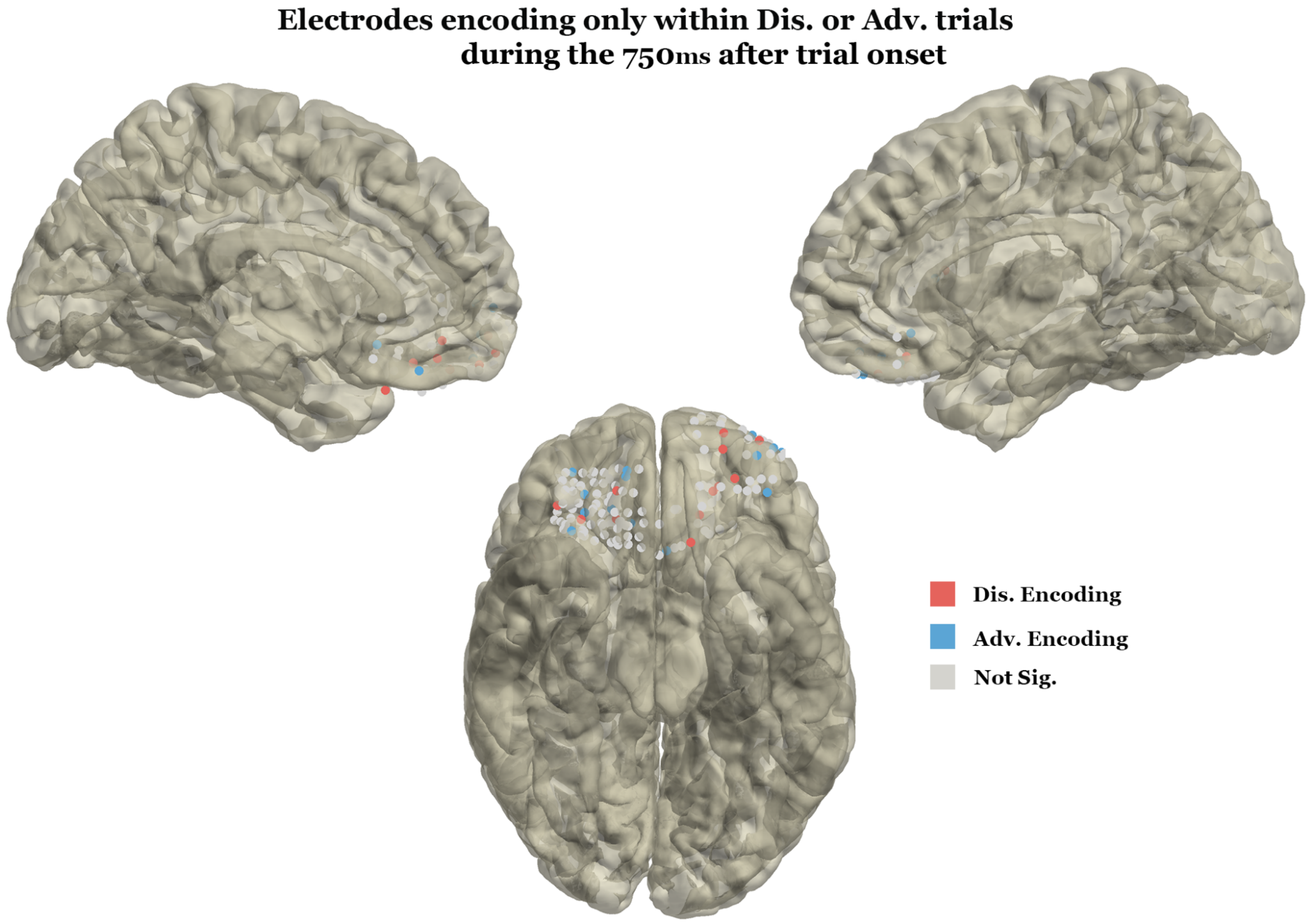


**Supplemental Figure 3.** Anatomical localization of state-dependent inequity encoding electrodes in the presentation epoch (blue/red/gray dots represent electrodes encoding only in advantageous/disadvantageous/none conditions).

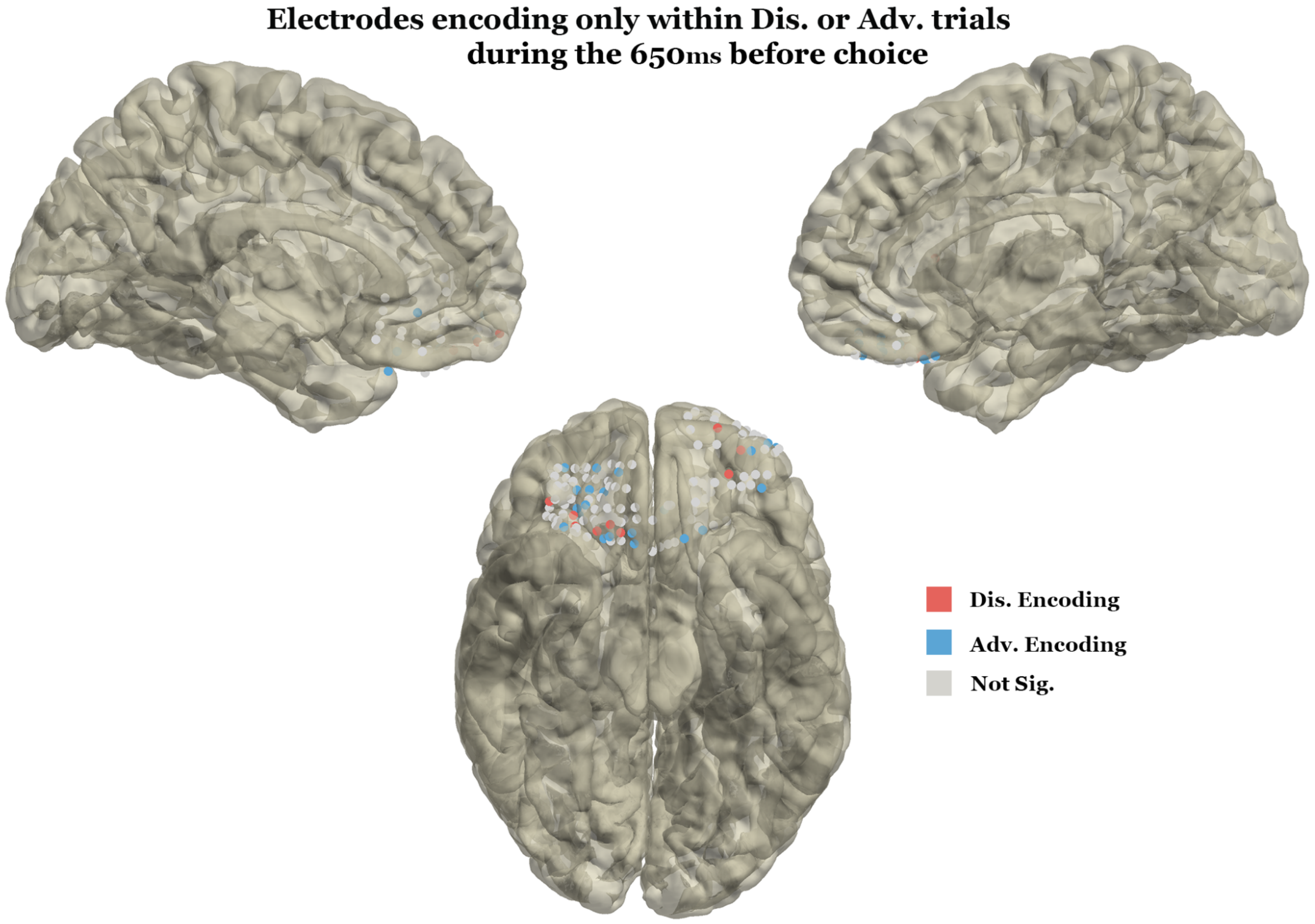


**Supplemental Figure 4.** Anatomical localization of state-dependent inequity encoding electrodes in the pre-choice epoch (blue/red/gray dots represent electrodes encoding only in advantageous/disadvantageous/none conditions).
